# Impact of a ketogenic diet intervention during radiotherapy on body composition: III. An interim analysis of the KETOCOMP study

**DOI:** 10.1101/343251

**Authors:** Rainer J. Klement, Gabriele Schäfert, Reinhart A. Sweeney

**Author notes:** Corresponding author Rainer Johannes Klement, Strahlentherapie Schweinfurt, Robert-Koch-Straße 10, 97422 Schweinfurt, Phone: 0049 / 9721 7202761.

## Abstract

**Background:** Ketogenic therapy (KT) in the form of ketogenic diets (KDs) and/or supplements that induce nutritional ketosis have gained interest as a complementary treatment for cancer patients. Besides putative anti-tumor effects, preclinical and preliminary clinical data indicate that KT could induce favorable changes in body composition of the tumor bearing host. Here we present first results of our ongoing KETOCOMP study (NCT02516501) study concerning body composition changes among rectal, breast and head & neck cancer (HNC) patients who underwent concurrent KT during standard-of-care radiotherapy (RT).

**Methods:** Eligible patients were assigned to one of three groups: (i) a standard diet group; (ii) a ketogenic breakfast group taking 50-250 ml of a medium-chain triglyceride (MCT) drink plus 10 g essential amino acids in the morning of RT days; (iii) a complete KD group supplemented with 10 g essential amino acids on RT days. Body composition was to be measured prior to and weekly during RT using 8-electrode bioimpedance analysis. Longitudinal data were analyzed using mixed effects linear regression.

**Results:** A total of 17 patients underwent KT during RT thus far (rectal cancer: n=6; HNC: n=6; breast cancer: n=5). All patients consuming a KD (n=14) reached nutritional ketosis and finished the study protocol with only minor problems reported. Compared to control subjects, the ketogenic intervention in rectal and breast cancer patients was significantly associated with a decline in fat mass over time (−0.3 and −0.5 kg/week, respectively), with no significant changes in skeletal muscle mass. In HNC patients, concurrent chemotherapy was the strongest predictor of body weight, fat free and skeletal muscle mass decline during radiotherapy, while KT showed significant opposite associations. Rectal cancer patients who underwent KT during neoadjuvant RT had significantly better tumor response at the time of surgery as assessed by the Dworak regression grade (median 3 versus 2, p=0.04483).

**Conclusions:** While sample sizes are still small our results already indicate some significant favorable effects of KT on body composition. These as well as a putative radiosensitizing effect on rectal tumor cells need to be confirmed once the final analysis of our study becomes possible.

## Introduction

Cancer patients frequently seek additional possibilities to support their standard therapies, improve their quality of life and positively influence their outcomes. One such supportive treatment approach is ketogenic therapy (KT) which comprises dietary interventions leading to nutritional ketosis such as ketogenic diets (KDs), short-term fasting and ketone body supplementation [1–3]. Nutritional ketosis is a physiological state, usually defined as β-hydroxybutyrate (BHB) concentrations exceeding 0.5 mmol/l [4], Ketogenic therapy for cancer patients is an emerging research topic, paralleled by an increasing interest on behalf of patients. For example, a survey among high grade glioma patients revealed that almost three quarters (73%) of patients would be willing to try a KD for three months and 66% would participate in a clinical trial investigating the effectiveness of the KD [5].

In a variety of preclinical tumor models, ketogenic therapy has shown beneficial effects, including efficacy against tumor growth and a positive impact on body composition, although some counterexamples showing no or even tumor-promoting effects of KDs or ketone bodies exist [6–9]. These contrasting findings concerning the efficacy of ketogenic therapy against tumor growth are most likely explained by the metabolic phenotype of the particular tumor treated [10–12], However, a growing number of studies reveal synergistic effects of ketogenic therapy with other therapies inducing oxidative stress in tumor cells such as radiotherapy (RT), chemotherapy or hyperbaric oxygen [1,3,13–15]. In addition, mechanistic studies provide evidence for muscle-sparing effects of ketone bodies, especially under conditions of insulin resistance often encountered in cancer patients. This makes sense from an evolutionary perspective, given that ketosis during starvation periods could have helped to maintain muscle mass which is indispensable for hunting and gathering foods.

Despite the growing number of preclinical in vitro and in vivo studies, research on the effects of ketogenic therapy in humans is still limited to small pilot studies and case reports [8]. In an initial case series of patients undertaking a KD during RT in our clinic we found some evidence that the diet could induce beneficial effects on body composition and quality of life [16]. This lead to the initiation of a clinical phase I study with the main aim to investigate the impact of a KD intervention on body composition in cancer patients undergoing RT (the KETOCOMP study, ClinicalTrials.gov Identifier: NCT02516501) [17]. Here we report an interim analysis of in total 63 patients who were either enrolled in the KETOCOMP study or followed a very similar protocol prior to study initiation. While the study is ongoing, these results are useful for providing first insights into the feasibility and effects of ketogenic therapy during RT treatment of ambulatory patients.

## Materials and Methods

### Study protocol

The KETOCOM P study has been approved by the ethics committee of the Bavarian Medical Association (Landesarztekammer Bayern) and registered under ClinicalTrials.gov Identifier no. NCT00123456. The detailed study protocol has been published previously [17]. Briefly, patients between 18 and 75 years with rectal, breast or head and neck cancer (HNC) referred to our clinic for curative RT were principally eligible for participating. Exclusion criteria were body mass index <18 or >35 kg/m^2^, Karnofsky index <70, pregnancy, metallic body parts that interfere with bioimpedance analysis (BIA), type I diabetes, known enzyme defects that contradict a KD and renal insufficiency. Patients were assigned to one of three groups: (i) a standard diet group; (ii) a ketogenic breakfast group taking 50-250 ml of a medium-chain triglyceride (MCT) drink (betaquick^®^, vitaflo, Bad Homburg, Germany) plus 10 g essential amino acids (MyAmino^®^, dr. reinwald healthcare gmbh & co kg, Altdorf, Germany) in the morning of RT days; (iii) a complete ketogenic diet group supplemented with 10 g essential amino acids on RT days. The composition of the MCT drink and amino acid supplement is shown in Supplementary Table 1.

Body composition was supposed to be measured prior to and weekly during RT using the seca mBCA scale (seca Deutschland, Hamburg, Germany). Based on body weight (BW), height, age, gender and 5 kHz und 50 kHz resistance and reactance values, the scale estimates fat free mass (FFM), extracellular water and total body water and - using 50 kHz values only - skeletal muscle mass (SMM) [18,19]. Fat mass (FM) was calculated as FM=BW-FFM. In order to standardize each measurement, patients were advised to fast overnight, not to drink in the morning and to void their bladder; their RT appointments were accordingly scheduled in the morning so that they could receive radiotherapy after BIA and weighing. On three occasions, blood samples were supposed to be collected with the patient still in fasting state immediately following BIA: once prior to, once in the middle of and once in the last week of RT.

As a proxy for measuring possible synergistic effects between ketogenic therapy and RT, we used the Dworak regression grade [20] at the time of surgery in rectal cancer patients treated with neoadjuvant chemo-RT. The Dworak grade ranges from 0 to 4, with 0 indicating no response and 4 complete remission of the tumor.

### Ketogenic interventions

In most cases, ketogenic interventions were started following baseline measurements prior to the first RT fraction and lasted until the final week of RT. Patients in the KETOCOMP study received a popular book on the KD for cancer patients [21], handouts with brief descriptions which foods to consume and which to avoid, urinary ketone strips for self-assessment of ketosis, and they had the opportunity to speak to our dietician. The consumption of a whole food KD was promoted, with emphasis on high-quality protein (meat, eggs and fish), micronutrient-dense foods (vegetables to every meal, organ meats and bone broth), and avoidance of industrial and processed foods (with the exception of MCT oil) as well as vegetable oils (except virgin coconut and olive oil) and foods rich in anti-nutrients (grains and legumes). Dairy products were suggested in moderation and preferably in the form of butter, cheese and fermented products. Due to the theoretically high micronutrient density and the brief duration of the KD, no additional supplements were advised. Patients in the ketogenic breakfast group were informed about the nature of a ketogenic diet and advised to avoid sugar and processed carbohydrates. They were instructed to receive each RT fasted and then consume the breakfast which they received by the technical staff. For the rest of the day, they could eat and drink ad libitum. Patients in the control group received no dietary advice, but were also free to receive dietary counseling, in which case they obtained the official recommendations of the German Society for Nutrition (DGE). Besides diet, all patients were advised to maintain their habitual lifestyle habits during the duration of RT.

### Study cohort

A list of patients included in this analysis is given in Supplementary Table 2, and from now on individual patients will be referred to by their number given in that table. In order to increase patient number and therefore modeling accuracy, a total of five patients undertaking a KD during RT prior to the KETOCOMP study initiation were included in this analysis since they received the same weekly BIA measurements as the KETOCOMP participants; four of these patients are described in more detail in a previous publication [16]. We also included one HNC patient who wished to participate in the KD group of the study despite having a metallic knee implant and included her data in the analysis of body weight changes. Due to the small number of patients recruited for the ketogenic breakfast group to date, patients receiving a ketogenic breakfast and those on a complete ketogenic diet were grouped together into a single “Ketogenic therapy” (KT) group.

### Statistical analysis

Longitudinal body composition data were analyzed using linear mixed effects models with the intercept and slope for time since start of RT as random effects depending on the individual patient. Time, intervention group (O=control/l=ketogenic), their interaction and the corresponding baseline body composition measure were included into each model. In addition, the following covariates were included based on their possible influence on body composition: Age, gender, baseline BMI, irradiated volume (planning target volume), and, for HNC patients, chemotherapy (0=no/1=yes) and PEG use in the timeframe prior to a particular measurement. For HNC patients, a time × chemotherapy and time × PEG use interaction were included if Akaike’s information criterion indicated an improvement in model fit. To ease interpretability of the regression coefficients, prior to model fitting, the covariates age and BMI were scaled to have mean zero and standard deviation 10 years or 10 kg/m^2^, respectively.

Differences between continuous and categorical variables were assessed using the Wilcoxon rank sum and Fisher’s exact test, respectively. All analysis was carried out in R, version 3.4.1 with the software package lme4 for linear mixed effects modeling.

## Results

Patient characteristics at baseline are given in Table 1. The KT and control groups were comparable with respect to most variables except for significantly lower BMI in the rectal and HNC intervention groups and higher fasting blood glucose levels in the rectal cancer control group. Minor deviations from the study protocol were the inclusion of a 76 year old rectal cancer patient and three breast cancer patients having BMI > 35 kg/m^2^ (maximum 36.57 kg/m^2^). Also, some patients received baseline measurements after RT had already started, but all within the first week of RT. The median study duration was 35 (KT) versus 37 (control) days (p=0.118) in breast cancer patients, 41 versus 43 days (p=0.9669) in HNC patients and 39 versus 34 days in rectal cancer patients (p=0.0277).

**Table 1:** Baseline characteristics of all patients.

|  | Rectal cancer |  |  | Head & neck cancer |  |  | Breast cancer |  |  |
| --- | --- | --- | --- | --- | --- | --- | --- | --- | --- |
|  | Ketogenic therapy (n=6) | Control (n=20) | p-value | Ketogenic therapy (n=6) | Control (n=14) | p-value | Ketogenic therapy | Control | p-value |
| Age [years] | 59 (38-74) | 65 (43-76) | 0.3141 | 63 (58-68) | 64 (55-75) | 0.8037 | 50 (42-72) | 58 (41-68) | 0.826 |
| Gender | Male: 4<br>Female: 2 | Male: 12<br>Female: 8 | 1 | Male: 5<br>Female: 1 | Male: 10<br>Female: 4 | 1 | Female: 5 | Female: 8 | 1 |
| BMI [kg/m<sup>2</sup>] | 24.4 (20.7-30.2) | 27.5 (19.5-32.8) | 0.03125 | 20.7 (19.3-26.2) | 24.4 (18.0-35.6) | 0.04076 | 28.7 (21.4-36.0) | 26.8 (18.8-36.6) | 0.8329 |
| Fasting glucose [mg/dl] | 92 (84-99) | 100 (66-265) | 0.01363 | 107 (98-154) | 107 (83-188) | 0.9671 | 91 (82-114) | 94 (81-113) | 0.7336 |
| Fasting BHB [mmol/l] | 0.15 (0.03-0.81) | 0.12 (0.05-0.6) | 1 | 0.23 (0.07-0.9) | 0.11 (0.03-0.42) | 0.1604 | 0.12 (0.03-0.27) | 0.11 (0.03-0.29) | 0.670 |
| 50 kHz phase angle [°] | 4.94 (4.74-6.59) | 4.69 (3.31-5.57) | 0.1103 | 4.59 (4.22-4.96) | 4.53 (3.96-5.70) | 0.9262 | 4.42 (4.22-5.39) | 4.47 (3.83-5.27) | 0.5237 |
| PTV [ccm] | 1402 (1081-1845) | 1473 (1098-2078) | 0.5727 | 805 (265-1278) | 816 (132-1147) | 0.7791 | 1116(560-1296) | 1102 (638-1658) | 0.7242 |
| Chemotherapy | Yes: 6 | No: 1<br>Yes:19 | 1 | No: 1<br>Yes: 5 | No: 5<br>Yes: 9 | 0.6126 | No: 5 | No: 8 | 1 |

Continuous and categorical variables were compared using the Wilcoxon rank sum and Fisher's exact test, respectively.

### Ketogenic intervention

#### Ketogenic diets

A total of 14 patients had undertaken a KD during RT: five rectal cancer, four HNC and five breast cancer patients (Supplementary Table 1). All of them exhibited nutritional ketosis with at least one finger prick or laboratory BHB concentration measurement reading >0.5 mmol/l. Median BHB concentrations during the KD were 0.6 (range 0.05-4.15) mmol/l, significantly higher than baseline values (p=0.0001857) and values measured in the controls (p=1.355×l0^−15^). All patients on a KD maintained compliance until the final measurement which marked the end of the study, except for one patient (#4 in Supplementary Table 2) who interrupted the diet for a few days due to aversion against fatty foods (especially the taste of coconut oil). Most patients also received 10 g of the essential amino acid supplement on radiation days which was tolerated by all but one who ingested the amino acids dissolved in water via a PEG tube (patient #29).

#### Ketogenic breakfast

So far, two patients maintained the maximum MCT dose during RT: Patient #7 revealed BHB concentrations of 1.1 mmol/l one hour after ingesting 50 g MCT fat, and patient #35 had 0.7 mmol/l ketosis 45 min after ingesting 45 g MCT fat. The lower maximum dose in the latter patient was due to a new packaging of the betaquick^®^ drink which reduced the volume of one container from 250 ml to 225 ml. One head and neck cancer patient (#33) had to reduce the MCT dose after reaching the maximum dose due to gastrointestinal problems and continued with about 20 g MCT fat per day which he took at home since he preferred a calm environment. He continued the study protocol with the amino acid supplement, added a ketogenic formula drink (KetoDrink, Tavarlin, Darmstadt, Germany) and additionally fasted a minimum of 16 h prior to each chemotherapy cycle. His median fasting BHB concentration during RT was 0.6 (0.45-0.82) mmol/l, and he accordingly was classified into the KT group. Two patients (#6 and #34) showed poor compliance to the ketogenic breakfast protocol in general and specifically complained about the taste of the amino acid supplement and gastrointestinal problems after consuming the MCT drink; they dropped out of the study early and were excluded from further analysis.

### Body composition changes

On average, 7 BIA measurements were performed per patient (range 2-9). Figure 1 shows linear regression lines for each patient stratified according to intervention group and tumor entity. Visually it appears that linear regression against time gives an adequate fit to the data; indeed, including a quadratic time component into the body composition models did not yield better fits as judged by the AIC. In fitting all the data for each tumor entity together, we found mixed effects models with varying slope and intercept superior to varying intercept only or fixed effects models as judged by both the AIC and maximum likelihood ratio test (results not shown). The results are given in Tables 2–4 for rectal, HNC and breast cancer patients, respectively.

**Figure 1:**
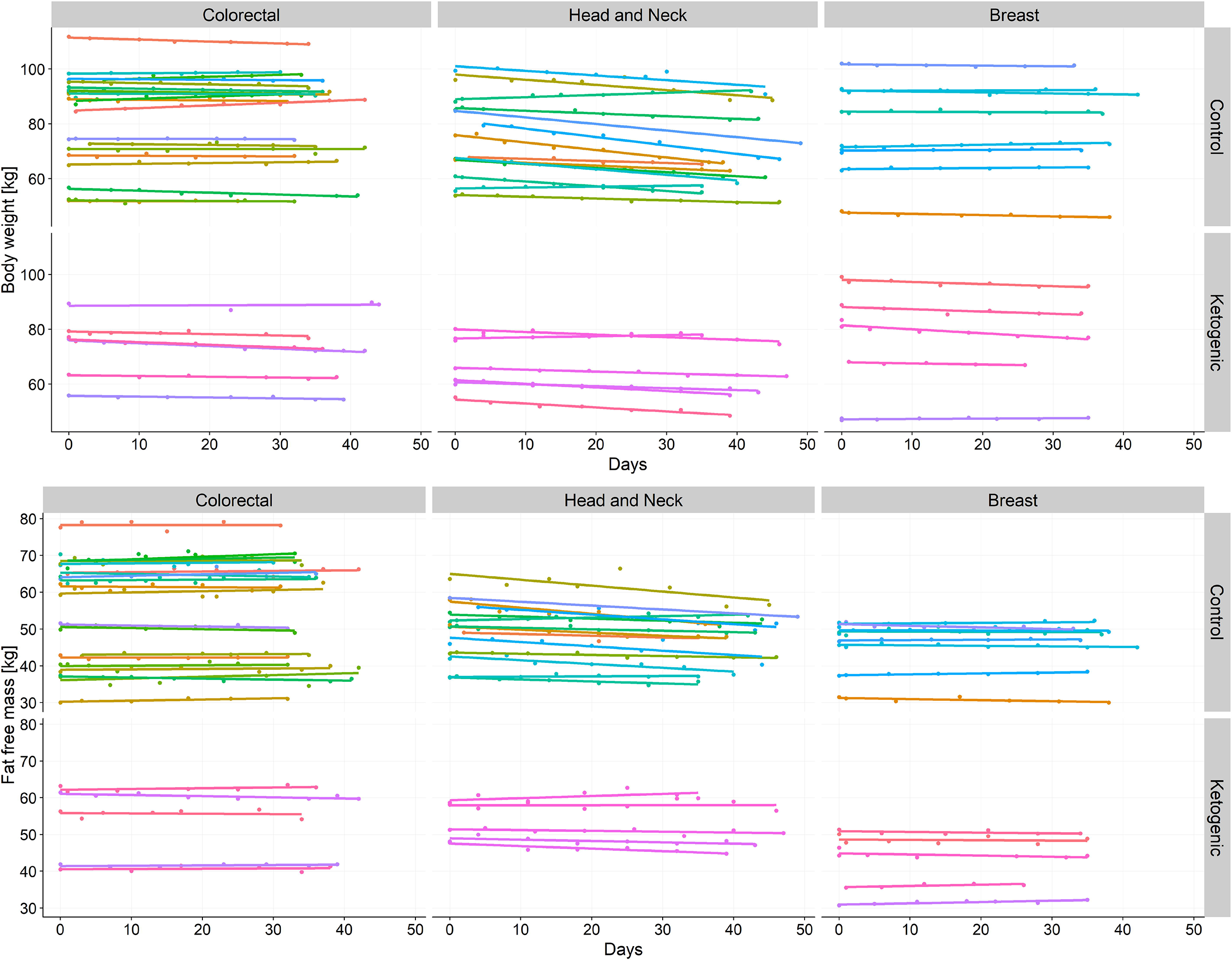
Changes in body weight and fat free mass during radiotherapy, stratified according to tumor site and intervention group.

**Table 2:**
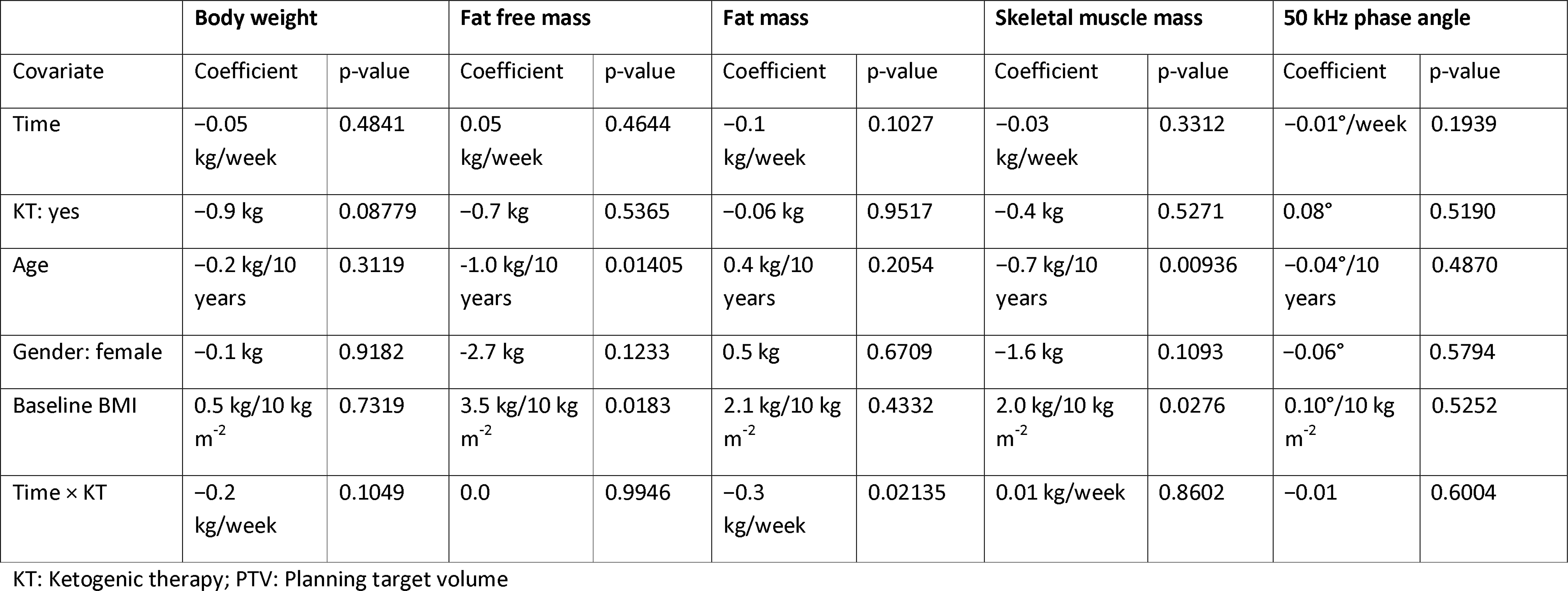
Regression coefficients for body composition changes in rectal cancer patients.

**Table 3:** Regression coefficients for body composition changes in head and neck cancer patients.

|  | Body weight |  | Fat free mass |  | Fat mass |  | Skeletal muscle mass |  | 50 kHz phase angle |  |
| --- | --- | --- | --- | --- | --- | --- | --- | --- | --- | --- |
| Covariate | Coefficient | p-value | Coefficient | p-value | Coefficient | p-value | Coefficient | p-value | Coefficient | p-value |
| Time | -0.3 kg/week | 0.09782 | -0.1 kg/week | 0.2714 | -0.2 kg/week | 0.2026 | -0.1 kg/week | 0.2073 | 0.01°/week | 0.6324 |
| KT: yes | -0.3 kg | 0.6338 | 0.5 kg | 0.3553 | -0.8 kg | 0.01466 | -0.04 kg | 0.8884 | -0.29° | 0.1431 |
| Age | 0.5 kg/10 years | 0.3423 | 0.5 kg/10 years | 0.2844 | -0.6 kg/10 years | 0.08465 | 0.2 kg/10 years | 0.3596 | -0.21°/10 years | 0.2154 |
| Gender: female | -1.0 kg | 0.1603 | 0.1 kg | 0.8950 | -0.02 kg | 0.9695 | 0.6 kg | 0.2369 | -0.03° | 0.8999 |
| Baseline BMI | 4.6 kg/10 kg m<sup>-2</sup> | 0.01475 | 1.1 kg/9.5 kg m<sup>-2</sup> | 0.06917 | 1.2 kg/10 kg m<sup>-2</sup> | 0.2340 | 0.2 kg/10 kg m<sup>-2</sup> | 0.4794 | -0.28°/10 kg m<sup>-2</sup> | 0.0900 |
| PEG use: yes | -0.1 kg | 0.6849 | -0.4 kg | 0.4279 | -0.1 kg | 0.6849 | 0.1 kg | 0.5675 | 0.22° | 0.07496 |
| Chemotherapy: yes | -0.7 kg | 0.2594 | 0.0 kg | 0.9950 | 0.01 kg | 0.9810 | 0.3 kg | 0.2210 | 0.26° | 0.2026 |
| Time × KT | 0.8 kg/week | 0.000148 | 0.6 kg/week | 0.0004163 | 0.3 | 0.1058 | 0.3 kg/week | 7.873×10<sup>-5</sup> | 0.005°/week | 0.8574 |
| Time × Chemotherapy | -1.2 kg/week | 1.655×10<sup>-8</sup> | -0.6 kg/week | 3.316×10<sup>-5</sup> | -0.5 | 0.0004188 | -0.5 kg/week | 2.361×10<sup>-9</sup> | -0.08°/week | 0.001794 |
| Time × PEG use | – | – | – | – | – | – | – | – | -0.02°/week | 0.5081 |

KT: Ketogenic therapy; PTV: Planning target volume

**Table 4:**
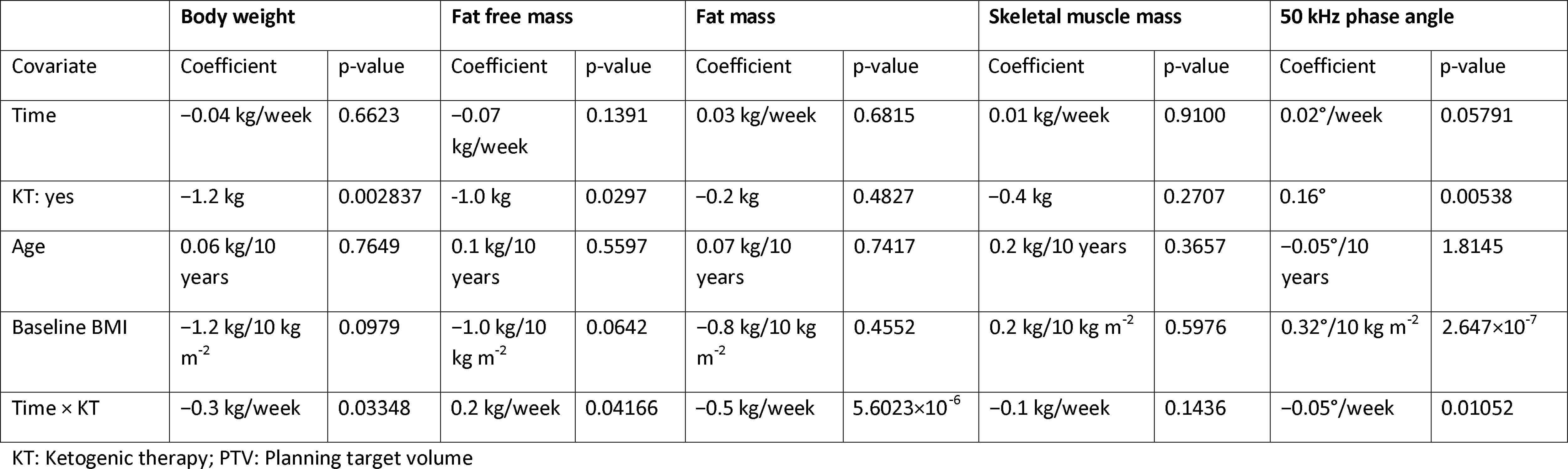
Regression coefficients for body composition changes in breast cancer patients.

#### Rectal cancer

In rectal cancer patients, those in the KT group lost significantly more BW between the first and the final measurement than those in the control group (ΔBW=2.05 kg versus 0.4 kg, p=0.04144). There was also a trend for a greater reduction in FM in the KT group (ΔFM= 1.7 kg versus 0.6 kg, p=0.09697). Changes in FFM, SSM and phase angle between first and final measurement were not significantly different between groups.

In linear regression analysis, KT was associated with a gradual loss of 0.2 kg BW and 0.3 kg FM per week, the latter association being significant (p=0.02135). No further significant associations with body composition were obtained (Table 2).

#### Head and neck cancer

Most HNC patients tended to lose some BW over the course of RT. Average weight loss was 5.0±4.3 kg in all patients and significantly greater in patients having received concurrent chemotherapy (7.0±2.9 kg versus 0.2±2.8 kg, p=0.0002064). Among patients having received chemotherapy, those in the KT group experienced significantly less weight loss than those in the control group (4.7±1.7 kg versus 8.3±2.7 kg, p=0.01199). In linear regression modeling, the strongest predictor of weight loss was chemotherapy (−1.2 kg/week), while KT was significantly associated with a weight gain of 0.8 kg/per week (Table 3). Chemotherapy was further strongly associated with gradual decreases in FFM, FM, SMM and phase angle, while KT had the opposite effect. In particular, KT significantly predicted for retention of FFM and SMM. Incorporation of a time × PEG use interaction did not improve the model fits except for phase angle where it was, however, not a significant predictor.

#### Breast cancer

All breast cancer patients in the KT group lost adipose mass between the first and the final measurement (median ΔFM=2.0 kg, range 1.3-4.2 kg) while there was basically no change in the control group control group (median ΔFM=0.07 kg, range −1.4-1.4 kg, p=0.003108 compared to the KT group). There was also a trend for greater BW loss in the KT group (ΔBW=3.1 kg versus 0.3 kg, p=0.06527). Changes in FFM, SMM or phase angle between the first and final measurement were not significantly different between groups.

In linear regression analysis, KT was associated with significantly greater gradual BW (−0.3 kg/week, p=0.03348) and FM (−0.5 kg/week, p=5.6023×l0^−6^) reductions (Table 4). While the model indicated a significant gradual decline in phase angle of 0.05°/week in the KT group, KT itself was associated with a 0.16° higher phase angle compared to the control group. The strongest predictor of phase angle was baseline BMI which indicated a 0.3° higher phase angle for every 10 kg/m^2^ increase in BMI.

### Tumor regression

Among the rectal cancer patients receiving neoadjuvant radio-chemotherapy (n=5 in the KT group, 19 in the control group), the median Dworak regression grade at the time of surgery was 3 (2-3) in the KT group compared to 2 (1-4) in the control group (p=0.04483), possibly indicating a better tumor response to RT. Patient #2 who was on a KD during RT refused surgery so that no Dworak grading was performed; he continued with the KD after RT and was progression-free at the last follow-up at 31 months after RT.

## Discussion

In this interim analysis of the ongoing KETOCOMP study, we investigated the effect of a ketogenic intervention - either a KD or a ketogenic breakfast containing highly bioavailable amino acids and ketogenic MCT fat - on body composition changes during RT. The majority of patients consumed a whole KD during the course of RT with no dropout. This is in stark contrast to some previous studies, especially the KETOLUNG and KETOPAN studies in which ≥50% of patients did not tolerate a highly artificial KD containing only 8% energy from protein during RT [22], We think the fact that our patients had early tumor stages, were intrinsically highly motivated and advised to eat a diet based on natural foods could have contributed to the good compliance. Our preliminary results indicate beneficial effects in terms of FFM and SMM retention in HNC patients and FM reduction in breast and rectal cancer patients. For rectal cancer, there was an indication of better tumor response to RT under the ketogenic intervention as assessed by the Dworak regression grade at surgery, although this result should be seen as preliminary due to the small number of patients.

It is increasingly recognized that BW per se is a poor indicator of nutritional status and health. BIA allows for an inexpensive, non-invasive tracking of body composition which has much more prognostic value since it is able to predict FFM, SMM and hydration status. By directly measuring the electrical properties of body tissues, BIA also provides additional clues about the nutritional status on the cellular level. For example, De Luis et al. showed that HNC patients were characterized by lower reactance and phase angle than healthy control subjects despite normal weight and BMI and even without prior weight loss [23]. On the metabolic side, these signs of cellular malnutrition manifest themselves as insulin resistance with increased lipid oxidation and impaired glucose tolerance [24–26]. Hence, it has been argued that high fat diets with an appropriate supply of amino acids provide the best metabolic support for the cancer patient while minimizing tumor growth promoting stimuli [27–29].

BIA is further useful to detect sarcopenia and sarcopenic obesity which are not straightforward to detect with standard anthropometric assessments, yet can have significant adverse consequences in terms of treatment tolerability and overall survival [30]. HNC patients represent a particularly frail population in this respect as they frequently develop sarcopenia during treatment which has been associated with poor quality of life and low physical performance status [31] and occurs even under recommended energy and protein intake [32], FFM loss can account for 60-70% of total weight loss in these patients and has been correlated to increased inflammatory cytokine and C-reactive protein levels [31,32]. It is therefore encouraging that our KT regime was associated with a significant retention of body weight, FFM and SMM during RT, directly opposing the effect of concurrent chemotherapy. Among weight losing HNC patients in our cohort (irrespective of concurrent chemotherapy) FFM accounted for 32.2% (29.4-54.1%) of body weight loss in the KT group and 48.7 (20.4-94.5%) in the control group (p=0.2121). If this gets confirmed with a larger number of patients it would support the hypothesis that ketosis with adequate high quality protein intake would protect against SMM loss in HNC patients [33].

In rats, ketosis has been shown to inhibit oxidation of the branched amino acids [34] and decrease the release of the gluconeogenic amino acid alanine [35]. Consistently, Sherwin et al. measured decreased nitrogen excretion and hypoalaninemia in fasting men upon BHB infusion while most other amino acid concentrations remained stable [36]. However, Phinney et al. detected significantly elevated branched chain amino acid levels in trained cyclists during a 4-week KD [37], Thus, in theory, nutritional ketosis could attenuate muscle protein catabolism while maintaining the availability of all amino acid precursors for muscle protein synthesis, leading to a net gain or at least maintenance of SMM despite lower insulin levels. Since it is availability of all essential amino acids that primarily drives muscle protein synthesis [38], the additional consumption of 10 g essential amino acids on radiation days could theoretically have further contributed to the attenuation of SMM loss in the HNC patients.

In rectal and breast cancer patients, FFM and SMM appeared to be maintained irrespective of the treatment group, although there was a significant association of KT with a gradual increase of FFM in breast cancer patients (Table 4). However, there was a significant correlation of KT with a gradual decline of FM in both patient populations and pronounced in breast cancer patients. This implies that KT would have increased the FFM-to-FM ratio in our patients, similar to our previous findings [16]. These findings are consistent with studies conducted in exercising individuals where short-term KDs resulted in FM loss while maintaining lean mass and performance [39–43]. Since most of our subjects did not exercise, a contribution of exercise-stimulated muscle protein synthesis can be ruled out as an explanation for the observed maintenance of SMM, and again the anti-catabolic effects of ketosis and/or anabolic effects of the supplemented essential amino acids may have contributed to the maintenance of SMM despite lower insulin levels and weight loss.

The reduction of FM can be rated as beneficial since adipose tissue has a putative role in promoting growth and survival of colorectal and breast cancer cells [44,45] and, accordingly, obesity has been found to be correlated with worse clinical outcomes in these patients [46–48]. Unfortunately, most breast cancer patients experience weight gain during therapy [49,50], and low carbohydrate diets have been proposed as an optimal countermeasure since they reduce insulin und blood glucose spikes, decrease adipose tissue, increase HDL cholesterol and decrease triglycerides and inflammation [51].

Further benefits of ketogenic diets have been found in preclinical studies in terms of direct antitumor effects that selectively sensitize tumor cells to radio-and chemotherapy [3,13–15]. While our data do not allow us to assess the putative anti-tumor effects of KT in terms of progression-free and overall survival, it is interesting that the rectal cancer patients undergoing KT had a significantly higher grade of tumor regression at the time of surgery than the control patients. However, because only five patients in the KT group received surgery, we interpret this result with caution until more data become available as the study proceeds. Nevertheless, our data would be consistent with antitumor effects of MCT-rich KDs in colon cancer animal models [52–54] while providing evidence against the hypothesis that ketone bodies could “fuel” rectal tumor growth as appears to be the case in a limited number of other animal studies [10,11]. Quite generally, the majority of animal studies thus far supports the hypothesis of anti-tumor effects of a KD [8]. Data supporting an anti-tumor action of KDs in patients with cancer of the colon or rectum are also emerging, including a case report of a rectal cancer patient on a paleolithic KD with 24 months follow-up [55] and a controlled clinical trial from Japan involving ten stage IV colon cancer patients treated with concurrent chemotherapy and a MCT-based KD for one year [56]. It is also interesting that Kato et al. [57] associated a KD-like eating pattern (defined as >40% energy from fat and <100 g/day glycemic load) with a reduced cancer-specific death risk of rectal cancer patients treated with RT.

The small number of patients under a ketogenic regime for each tumor entity poses one of the largest limitations of our results presented herein. However, with a median of 7 BIA measurements per patient we have collected enough data points for building mixed effects linear regression models with incorporation of several covariates with a putative influence on body composition. In the final analysis it is planned to separately evaluate patients in the ketogenic breakfast group and those eating a full KD.

Combining fully KD patients with those eating just a ketogenic breakfast into a single group poses another limitation of this analysis. Besides increasing the sample size, we justify this approach based on important similarities among both intervention groups including (i) the fact that at least a few hours after RT, both patient groups would have been in nutritional ketosis, (ii) that most patients from both groups took the same amount of the amino acid supplement, and (iii) that some of the HNC patients receiving the ketogenic breakfast also received additional ketogenic drinks for consumption at home. Unfortunately, it is not possible to separate the contributions of the amino acid supplement and ketosis to the observed beneficial effects on body composition. Nevertheless, we conceive the addition of crystalline essential amino acids to a KD regime as a good strategy to increase muscle protein synthesis without the need to increase the amount of food proteins which could interfere with ketosis.

Finally, the validity of BIA for estimating body composition is limited by assumptions relating to body shape. Comparing the estimates of our BIA device to those derived from Dual-energy X-ray Absorptiometry and MRI, Bosy-Westphal et al. calculated the coefficients of determination (R^2^) for the FFM and SMM prediction equations as 0.98 and 0.97, respectively, and the root mean square errors as only 1.9 kg and 1.2 kg, respectively [18,19]. Since we were mainly interested in changes of body composition and not their exact absolute values, our conclusions are robust against any systematic deviations from the true body composition values within individual patients.

## Conclusion

In this preliminary analysis we observed beneficial effects of a ketogenic intervention during RT on body composition: Rectal and especially breast cancer patients lost adipose tissue while preserving lean body mass and HNC patients lost significantly less BW, FFM and SMM compared to the control group. In addition, in five rectal cancer patients who combined KT and RT tumors had regressed more at the time of surgery than in control patients, a finding that reached statistical significance despite the small number of patients in the KT group. If the final data from the ongoing KETOCOMP study confirm these early results, it would provide a justification for using KT alongside RT for patients who are interested in taking self-responsibility to support their therapy.

## Acknowledgments

We thank Sabine Chwola for helping with coordinating the patient’s measurement appointments.

## Authorship statement

RJK and RAS designed the study and collected the data. RJK analyzed the data and wrote the initial manuscript draft. GS helped with conducting the dietary intervention. All authors read, edited and approved the final manuscript.

